# *Enterococcus faecalis* promotes innate immune suppression and polymicrobial catheter-associated urinary tract infection

**DOI:** 10.1101/133140

**Authors:** Brenda Yin Qi Tien, Hwee Mian Sharon Goh, Kelvin Kian Long Chong, Soumili Bhaduri-Tagore, Sarah Holec, Regine Dress, Florent Ginhoux, Molly A. Ingersoll, Rohan B. H. Williams, Kimberly A. Kline

## Abstract

*Enterococcus faecalis*, a member of the human gastrointestinal microbiota, is an opportunistic pathogen associated with hospital-acquired wound, bloodstream, and urinary tract infections. *E. faecalis* can subvert or evade immune-mediated clearance, although the mechanisms are poorly understood. In this study, we examined *E. faecalis*-mediated subversion of macrophage activation. We observed that *E. faecalis* actively prevents NF-κB signaling in mouse RAW264.7 macrophages in the presence of Toll-like receptor agonists and during polymicrobial infection with *Escherichia coli*. *E. faecalis* and *E. coli* co-infection in a mouse model of catheter-associated urinary tract infection (CAUTI) resulted in a suppressed macrophage transcriptional response in the bladder compared to *E. coli* infection alone. Finally, we demonstrated that co-inoculation of *E. faecalis* with *E. coli* into catheterized bladders significantly augmented *E. coli* CAUTI. Taken together, these results support that *E. faecalis* suppression of NF-κB-driven responses in macrophages promotes polymicrobial CAUTI pathogenesis.

**Author Summary:** Synergistic polymicrobial infections can contribute to both disease severity and persistence. *Enterococcus faecalis* and *Escherichia coli* are frequently co-isolated from polymicrobial urinary tract infections. Immunomodulation by co-infecting microbes can result in a more permissive environment for pathogens to establish infection. Presently, we do not yet understand how these microbes overcome host immunity to establish polymicrobial infections. To address this, we investigated how the immunosuppressive function of *E. faecalis* can contribute to acute infection. We defined that *E. faecalis* is able to suppress macrophages *in vitro*, despite the presence of *E. coli*. We also demonstrated *E. faecalis’* ability to augment *E. coli* titers *in vivo* to establish kidney infection. Our findings raise the prospect that *E. faecalis* can alter host immunity to increase susceptibility to other uropathogens.

## Introduction

*Enterococcus faecalis* is an early colonizer in infants and a ubiquitous member of the human gut microbiome [1]. *E. faecalis* is also associated with up to 70% of wound infections, nearly 10% of bloodstream infections, and up to 30% of catheter-associated urinary tract infections (CAUTI) [2-5]. To successfully colonize and persist in the host, pathogens must withstand, modulate, or evade immune-mediated clearance mechanisms. *E. faecalis* invokes multiple strategies to persist within the host, including the formation of biofilms that prevent phagocytosis by immune cells [6], and the ability to survive within macrophages and neutrophils for extended periods of time [7-11].

Mammalian cells detect pathogen-associated molecular patterns (PAMPs) via pattern recognition receptors (PRRs) to trigger nuclear factor-kappa B (NF-κB)-dependent host defenses. NF-κB controls the transcription of inflammatory and immune-associated genes, including cytokines and chemokines regulating recruitment and activation of immune cells in response to infection [12]. *E. faecalis* infection of macrophages at low multiplicities of infection (MOI = 10) results in the activation of mitogen activated protein kinases (MAPKs) and NF-κB, leading to the production of pro-inflammatory cytokines [13]. However, some *E. faecalis* strains, isolated from the gastrointestinal tract of healthy human infants, can suppress MAPK and NF-κB signaling, and IL-8 expression in intestinal epithelial cells *in vitro* [14, 15].

In a mouse urinary tract infection (UTI) model, the cellular response to *E. faecalis* infection is primarily monocytic and is independent of Toll-like receptor (TLR) 2 [16]. In a CAUTI model, the presence of a urinary catheter alone elicits a strong pro-inflammatory response in the bladder composed of neutrophils and monocyte-derived cells [17-19]. Infection of catheterized bladders with *E. faecalis* results in the development of high titer catheter-associated biofilms and bladder infection, despite the presence of the strong inflammatory response induced by catheterization (19). Moreover, in the course of *E. faecalis* CAUTI, the number of both non-activated and activated bladder-associated macrophages was decreased compared to catheterized, uninfected, animals [17]. Together, these observations suggest that *E. faecalis* can subvert immune-mediated killing to persist within the infected bladder.

During UTI and CAUTI, *E. faecalis* is often part of a polymicrobial community [20-22]. *E. faecalis* can promote polymicrobial infection by increasing the resistance of co-infecting organisms, such as *P. aeruginosa* and *P. mirabilis*, to clearance by antibiotics [23, 24]. Polymicrobial infection by *E. faecalis* and *P. aeruginosa* more often leads to aggravated pyelonephritis, compared to monomicrobial infection [23]. *E. faecalis* and uropathogenic *Escherichia coli* (UPEC) are also frequently isolated together during CAUTI [25], however, the relationship between these pathogens and the impact on pathogenesis is unknown. Given the frequency with which *E. faecalis* is found within polymicrobial infections, and that *E. faecalis* can modulate the host immune response within the catheterized bladder, we tested the hypothesis that *E. faecalis* immune modulation promotes polymicrobial CAUTI. We found that *E. faecalis* actively subverts *E. coli*-mediated NF-κB activation and pro-inflammatory cytokine production in RAW264.7 macrophages *in vitro* and macrophage-associated pro-inflammatory expression profiles in catheterized bladders *in vivo,* culminating in higher titer *E. coli* CAUTI.

## Results

### Live *E. faecalis* prevents LPS-or LTA-mediated NF-κB-driven activation in RAW macrophages

*E. faecalis* infection during CAUTI induces monocytic infiltration [16]. To determine whether *E. faecalis* immunomodulated monocyte-derived cells such as macrophages, we assessed NF-κB signaling in mouse RAW-264.7 macrophages at 6 hours post-infection (hpi). Both *E. faecalis* strain OG1RF **(Fig 1A)** and the multidrug-resistant strain V583 **(S1A Fig)** activated NF-κB at low multiplicities of infection (MOI) as previously reported [13]. By contrast, neither *E. faecalis* OG1RF nor V583 activated NF-κB signaling at high MOIs. We simultaneously monitored lactate dehydrogenase (LDH) release into culture supernatants to ensure that the absence of NF-κB activation was not a result of cell death at high MOIs, and observed no increase in LDH release at any of the MOIs used in this study **(Fig 1B and S1B Fig from S1 Fig)**.

**Fig 1.**
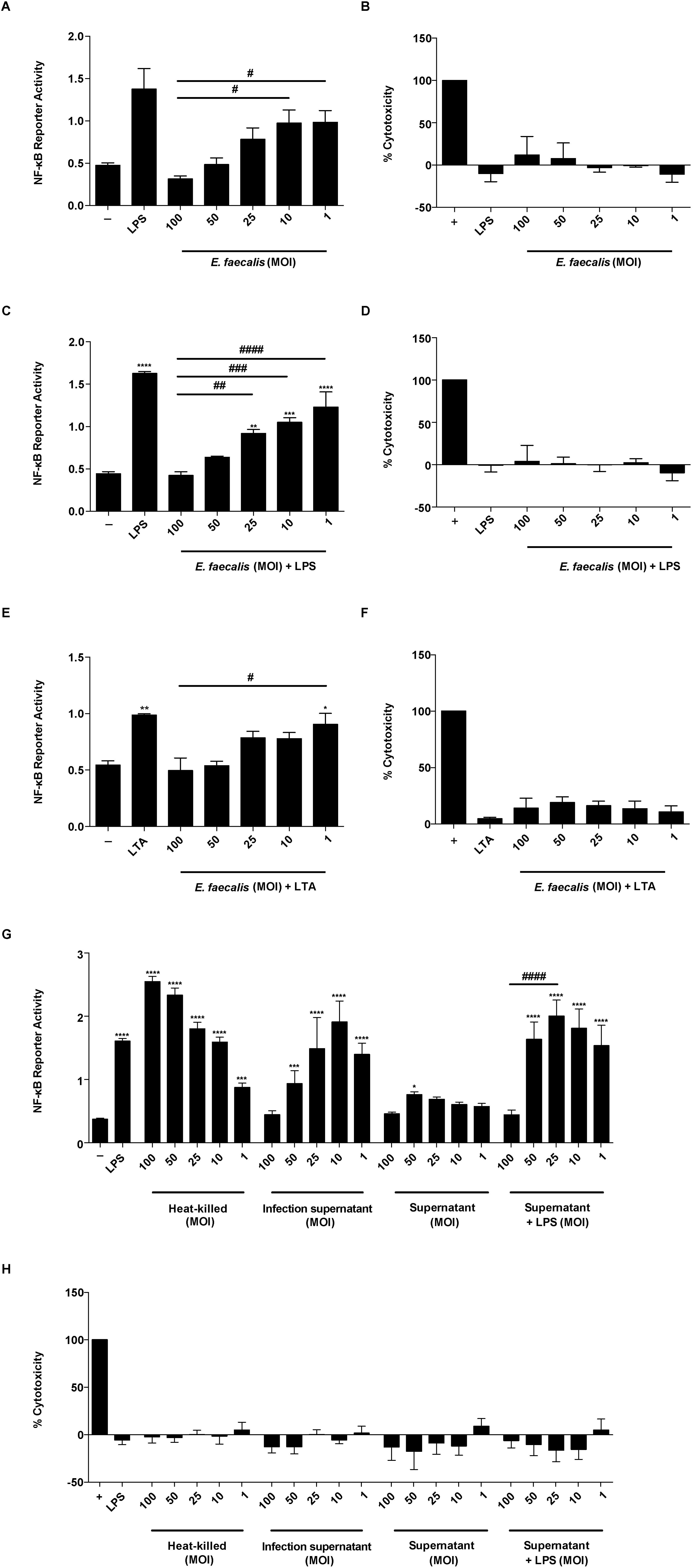
*E. faecalis* prevents NF-κB-driven macrophage activation. Mouse RAW 267.4 macrophages were infected with live *E. faecalis* OG1RF alone, or treated concurrently with either LPS (100 ng/ml) or LTA (100 ng/ml) at the specified MOI for 6 hours prior to measurement of NF-κB-driven SEAP reporter activity and cytotoxicity (LDH activity). (A) NF-κB-driven SEAP reporter activity and (B) LDH activity of RAW 264.7 macrophages infected by *E. faecalis* alone. (C) NF-κB-driven SEAP reporter activity and (D) LDH activity in the presence of *E. faecalis* and LPS. (E) NF-κB-driven SEAP reporter activity and (F) LDH activity in the presence of *E. faecalis* and LTA. (G) NF-κB-driven SEAP reporter activity and (H) LDH activity upon stimulation with heat-killed *E. faecalis* at the indicated MOI, infection supernatant, or bacterial culture supernatants with and without LPS. Culture supernatants post infection period were collected for SEAP reporter assays and LDH assays. NF-κB-driven SEAP reporter assays: exposure to media alone (-) represents background NF-κB reporter activity and stimulation with LPS or LTA represents positive controls for reporter activity. LDH assays: Triton-X treatment served as a positive control (+) for cell death. Data are combined from 3 independent experiments; mean values are graphed and error bars represent standard error of the mean (SEM). Statistical analysis was performed by the one-way ANOVA test with Tukey’s multiple comparison test where \**P*<0.05, \*\*\**P*<0.001, \*\*\*\**P*<0.0001 as compared to media alone (-) controls; and where ^#^*P*<0.05, ^##^*P*<0.01, ^###^*P*<0.001, ^####^*P*<0.0001 among all of the MOIs as compared to MOI 100.

*E. faecalis* can attenuate proinflammatory cytokine secretion in intestinal epithelial cells [15]. To determine whether *E. faecalis* actively prevented NF-κB-mediated transcription, or simply failed to induce NF-κB-mediated transcription at high MOIs in macrophages, we tested whether *E. faecalis* could prevent NF-κB-driven activation in the presence of TLR agonists that initiate NF-κB signaling. We exposed macrophages to lipopolysaccharide (LPS) or lipoteichoic acid (LTA) simultaneously with *E. faecalis* for 6 hours, quantified both NF-κB-mediated transcription and LDH release, and observed a dose-dependent inhibition of LPS-and LTA-induced NF-κB activation by *E. faecalis* **(Fig 1C, 1E and S1C Fig from S1 Fig)** in the absence of cytotoxicity **(Fig 1D, 1F and S1D Fig from S1 Fig)**.

To determine whether the absence of an NF-κB transcriptional response was due to an *E. faecalis* secreted factor, we examined the macrophage response to heat-killed *E. faecalis* or cell-free bacteria supernatants from MOI equivalents ranging from 100 to 1. We observed that heat-killed *E. faecalis* activated NF-κB at all MOIs, in an inverse manner to that of live intact cells **(Fig 1G)** in the absence of cytotoxicity **(Fig 1H)**. Supernatants from infected macrophages showed similar NF-κB activation to that of live *E. faecalis* cells **(Fig 1G)**. To rule out that NF-κB activation was not due to cytokines secreted by RAW-264.7 macrophages during infection, we also exposed macrophages to supernatants from bacteria cultures grown in the absence of macrophages. Supernatants from bacteria cultures weakly activated NF-κB alone and did not suppress LPS-mediated induction of NF-κB activity, except at MOI 100 **(Fig 1G)**. Together, these data suggest that *E. faecalis* actively prevented NF-κB activation via a process requiring a heat-modifiable factor that is secreted into cell supernatants during co-culture with macrophages, and that is produced in the absence of macrophages only at very high MOI.

### *E. faecalis* suppresses NF-κB-dependent cytokine and chemokine production in RAW macrophages

*E. faecalis* modulates cytokines such as IL-8, TNFα, and IL-1β in intestinal epithelial cells [13, 14]. To intestinal epithelial cells] To investigate whether *E. faecalis* suppresses cytokine production in infected macrophages, we measured release of a variety of cytokines and chemokines, whose expression is dependent on NF-κB activation, in the absence of LPS. We observed an overall increase of both pro-and anti-inflammatory cytokines and chemokines at MOI 10 and 1, similar to that observed in LPS-treated cells **(Fig 2A,B)**. Strikingly, at MOI 100, we observed a global decrease in cytokine, chemokine, and growth factor expression as compared to MOI 10 or LPS exposure **(Fig 2A)**. Moreover, at MOI 100, we observed that most of the analytes (IFN-□, CCL11, CSF2, IL-4, IL-17, IL-12p40, IL-12p70, IL-2, IL-1β, CCL2, CXCL1, and IL-5) were present at levels similar to the media control **(Fig 2A,B and S2A Fig from S2 Fig)**. Principal component analysis of analytes revealed that the profile of MOI 100 overlapped with the profile of uninfected macrophages, suggesting that analytes were not expressed despite greater numbers of *E. faecalis* **(S2B Fig from S2 Fig)**. Therefore, these data suggest that *E. faecalis* suppression of NF-κB activation at high MOI led to an overall suppression of cytokines and chemokine expression **(Fig 2)**.

**Fig 2.**
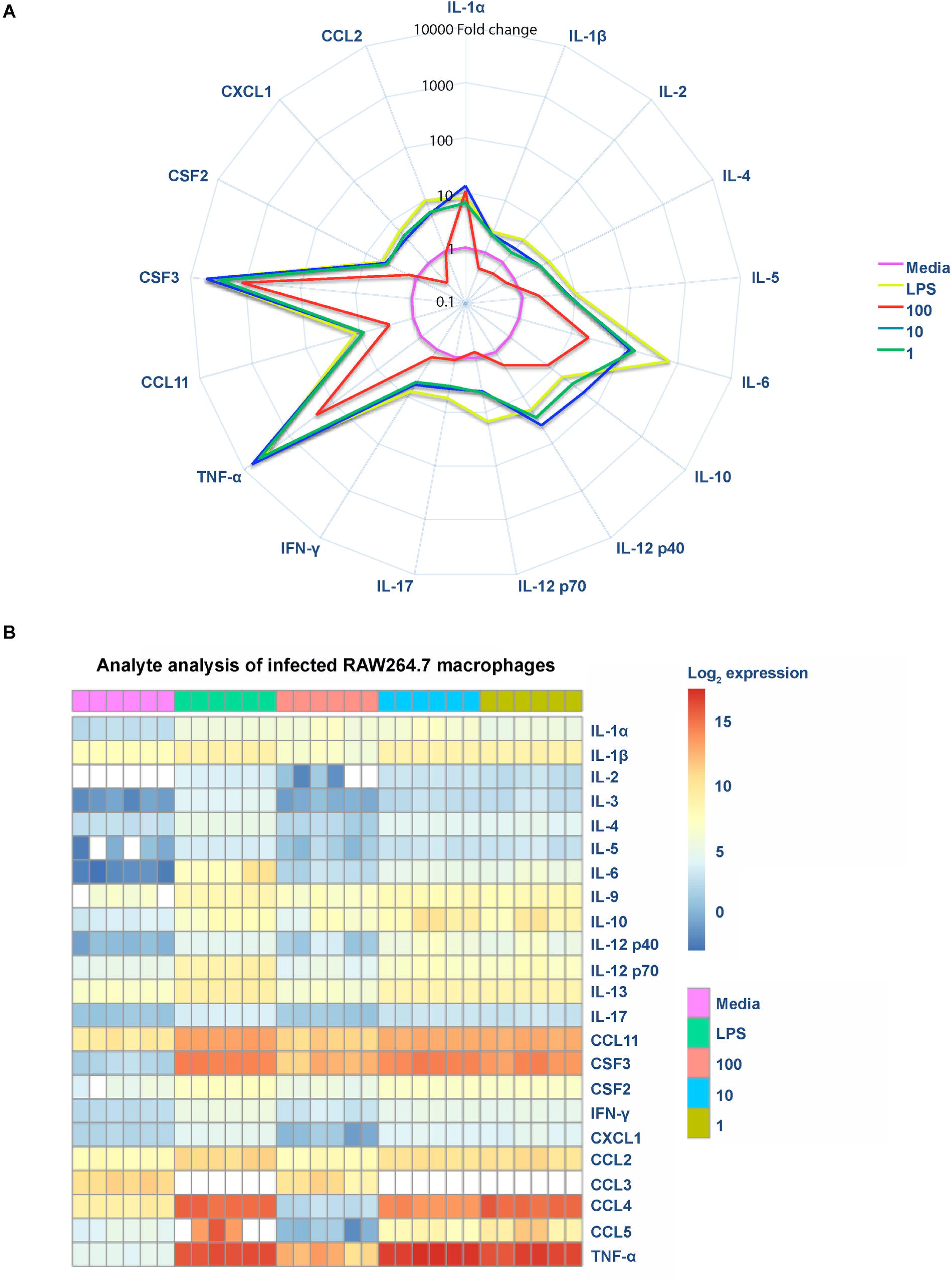
*E. faecalis* suppresses NF-κB-dependent cytokine and chemokine production in RAW macrophages. Mouse RAW 267.4 macrophages were infected with live *E. faecalis* at the indicated MOI. (A) Spider plot showing the fold-change of cytokines, chemokines and growth factors detected in filtered supernatants collected 6 hpi at depicted conditions compared to media control. Data were normalized against the media control, represented in pink, to obtain fold-change. (B) Heat map depicting the log_2_ transformation of absolute values measured in pg/ml of the indicated cytokines, chemokines, and growth factors.

### *E. faecalis* limits *E. coli*-mediated immune activation during polymicrobial RAW macrophage infection

To investigate whether *E. faecalis-*mediated immune suppression contributed to polymicrobial UTI, we first tested its ability to suppress NF-κB activity in the presence of *E. coli in vitro*. We determined that RAW macrophages infected with *E. coli* K12 strain MG1655 at an MOI of 1 or *E. coli* UTI89 at an MOI of 0.125 induced NF-κB activation **(S3A and S3C Fig from S3 Fig)** in the absence of cytotoxicity **(S3B and S3D Fig from S3 Fig)**, whereas higher MOIs were toxic to the mammalian cells **(S3B and S3D Fig from S3 Fig)**. We simultaneously infected macrophages with *E. faecalis* and *E. coli* at these pre-determined MOIs and observed that, while both *E. coli* strains MG1655 and UTI89 mono-infection induced NF-κB reporter activity equal to LPS alone, *E. faecalis* prevented *E. coli*-induced NF-κB activity in a dose-dependent manner **(Fig 3)**. From this observation, we hypothesized that *E. faecalis* could similarly suppress the host immune response *in vivo*.

**Fig 3.**
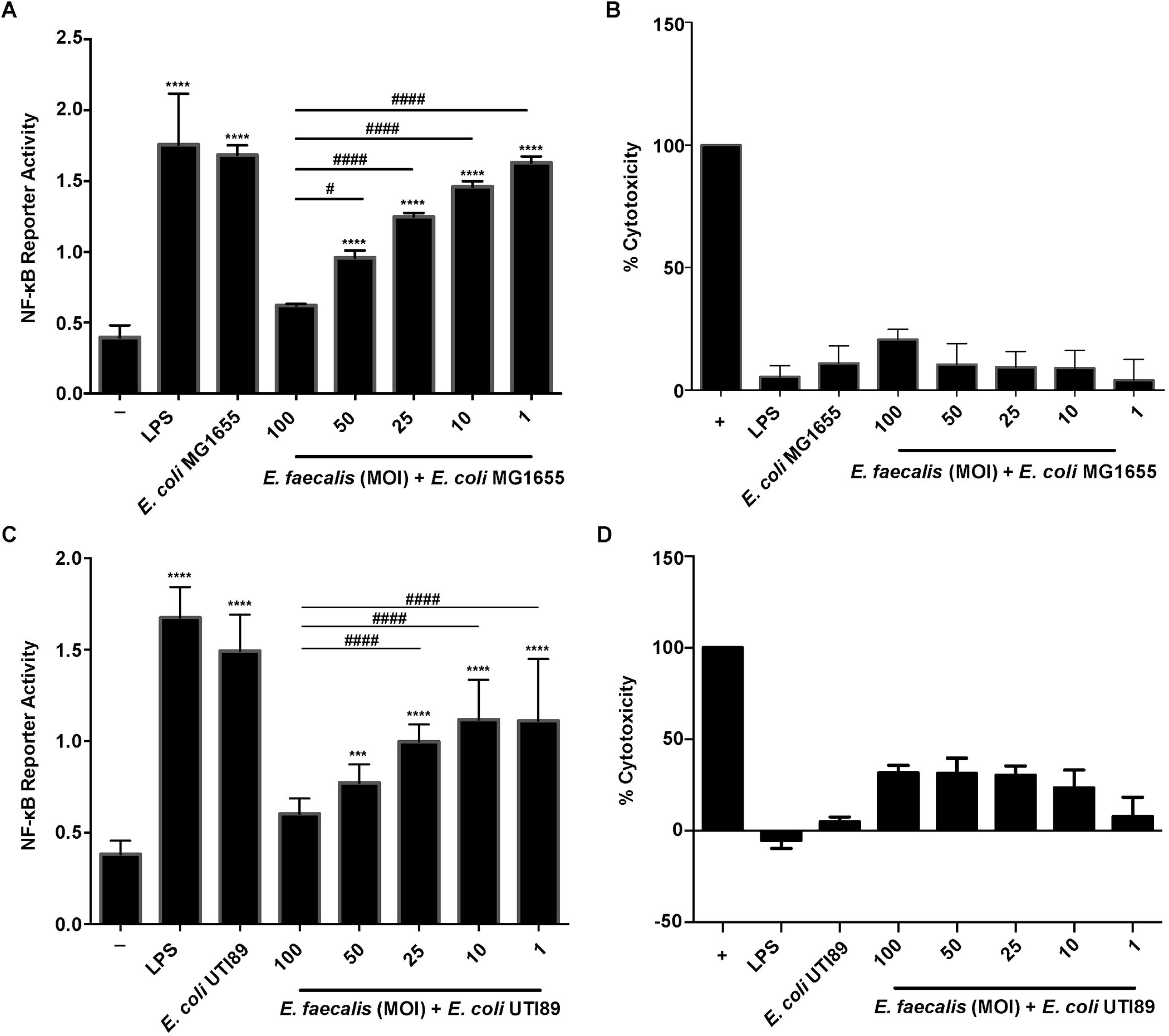
*E. faecalis* suppresses *E. coli* induced immune activation *in vitro*. Mouse RAW 267.4 macrophages were stimulated simultaneously with *E. faecalis* OG1RF and *E. coli* MG1655 prior to measurement of (A) NF-κB-driven SEAP reporter activity and (B) LDH activity. Mouse RAW 267.4 macrophages were co-infected with *E. faecalis* OG1RF and *E. coli* UTI89 before measuring (C) NF-κB-driven SEAP reporter activity and (D) LDH activity. NF-κB-driven SEAP reporter assays: exposure to media alone (-) represents background NF-κB reporter activity and stimulation with LPS represents positive controls for reporter activity. LDH assays: Triton-X treatment served as a positive control (+) for cell death. Data are combined from 3 independent experiments. Statistical analysis was performed using the one-way ANOVA test with Tukey’s multiple comparison test where \*\*\**P*<0.001, \*\*\*\**P*<0.0001 as compared to media alone (-) controls; and where ^#^*P*<0.05, ^###^*P*<0.001, ^####^*P*<0.0001 among all of the MOIs as compared to MOI 100.

### *E. faecalis* limits *E. coli*-mediated immune activation during mixed species infection

To investigate whether *E. faecalis* impacts immune-related signaling *in vivo*, we performed RNA expression profiling on whole bladders 24 hours post catheterization and infection. We compared *E. coli* UTI89 mono-species infection to *E. coli* UTI89 and *E. faecalis* OG1RF co-infection at a 1:1 inoculum ratio. Of the 15,501 detectable genes (*P*_*adj*_<0.05), 2 genes (0.013%) demonstrated increased mRNA levels, while 53 genes (0.34%) demonstrated decreased mRNA levels between co-infected mice and the mono-infected mice. Of these differentially expressed genes, we observed that 31 genes mapped to Gene Ontology (GO) terms: *response to external biotic stimulus* (GO:0043207), *response to other organism* (GO:0051707), *innate immune response* (GO:0045087), *response to cytokine* (GO:0034097), *response to biotic stimulus* (GO:0009607), *immune effector response* (GO:0002252) and *regulation of immune response* (GO:0050776), that were significantly enriched (*P*_*adj*_<0.01; Fisher’s Exact Test, corrected the 218 terms tested; see Methods section) **(Fig 4A and Fig 4B, Table from S1 Table).**

**Fig 4.**
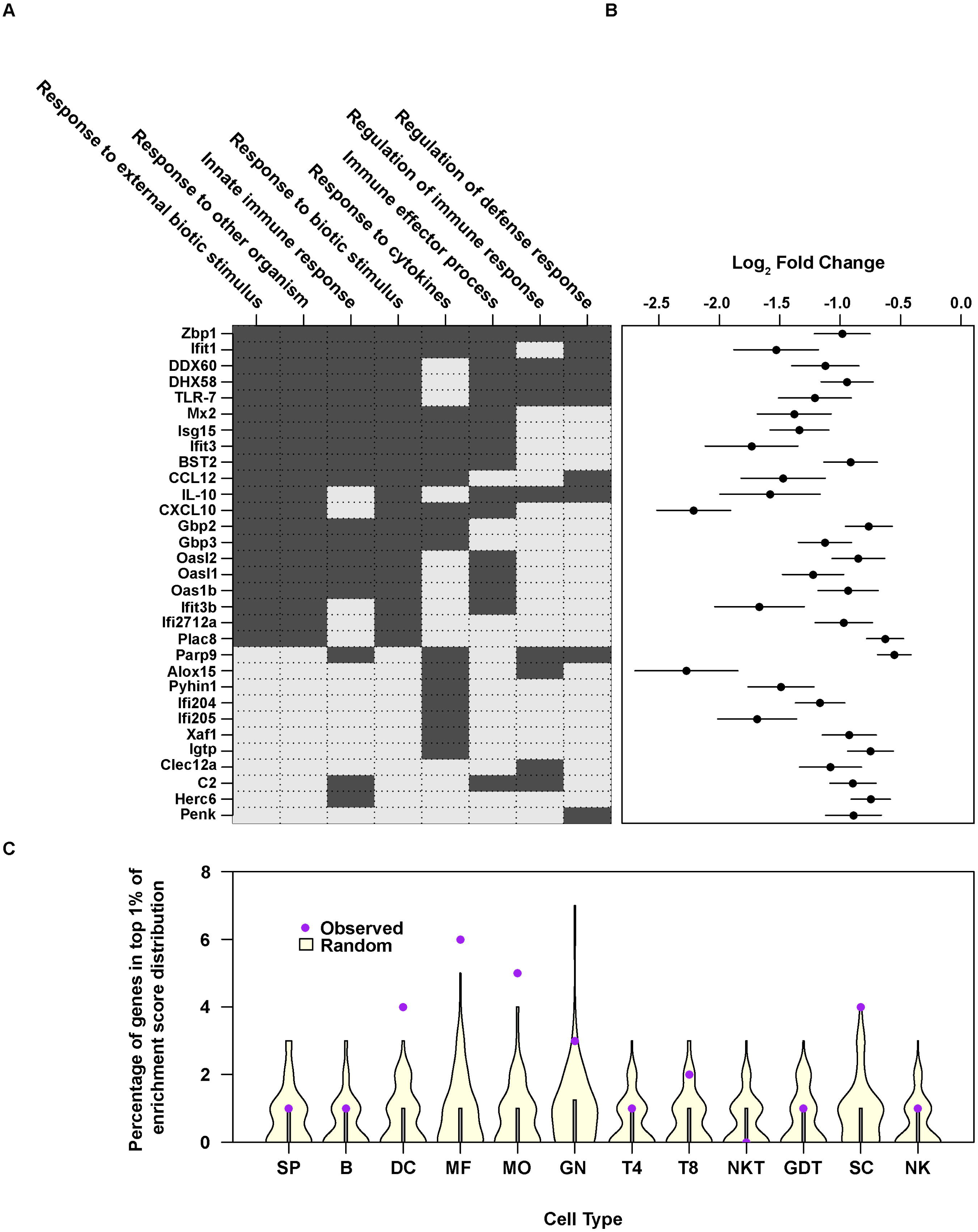
*E. faecalis* suppresses *E. coli*-driven inflammation in catheterized mouse bladders. Female C57BL/6NTac mice were implanted with catheters in the bladder and infected with 10^7^ CFU of *E. coli* UTI89 or 10^7^ CFU each of *E. coli* and *E. faecalis* in a 1:1 mixture. After 24 hours, bladders were removed and RNA extracted. (A) Binary matrix showing association between differentially expressed genes (rows) and those Gene Ontology Biological Process (GOBP) terms (columns) enriched in the differential expression analysis between *E. coli*-infected and polymicrobial-infected animals. Differentially expressed genes that did not map to an enriched GOBP term are not shown. Dark cells indicate genes that are annotated to a GOBP term and light cells indicate that it is not. (B) Summary of differential expression in set of 31 genes shown in (A). Differential expression in each gene is summarized by mean (black dots) log_2_ ratio of expression between *E. coli*-infected and polymicrobial-infected animals; where the line indicates estimated standard error of mean. (C) To examine whether observed differential gene expression may be associated with a given ImmGen defined cell type, we calculated the percentage of the top differentially genes (up-regulated in *E. coli* infected compared to polymicrobial infection) that are placed within the top 1% of the distribution of the cell-type-specific enrichment score [66] (purple dots) compared to 100 sets of 50 genes drawn at random (summarized in violin plots in yellow; the bar shows the median). SP, stem and progenitor cells; B, B cells; DC, dendritic cells; MF, macrophages; MO, monocytes; GN, granulocytes; T4, CD4+ cells; T8, CD8+ cells; NKT, natural killer T cells; GDT, γδ T cells; SC, stromal cells; NK, natural killer cells. See Tables S1, S2 for related analyses. The experiment was performed twice (n=3 mice per group/experiment). Representative data are shown from one experiment.

The enrichment of GO terms associated with immune function within down-regulated genes during co-infection in the presence *E. faecalis* suggested that we might also observe differential gene expression specifically in genes associated with the cell populations responding to CAUTI [19]. To test this, we examined the Immunological Genome Project (ImmGen) database, which comprises publicly available data from a collection of immune cell types in C57BL/6J mice [13, 26]. We found that within the top 50 differentially regulated genes between the mono-infected and co-infected groups, genes specific for dendritic cells (DC), macrophages (MF), and monocytes (MO) were over-represented and showed decreased mRNA levels in co-infected animals, suggesting a reduced infiltration or activation of these cells in the bladder following-co-infection as compared to mono-infection **(Fig 4C, Table from S2 Table)**.

### *E. faecalis* limits *E. coli*-mediated immune activation and promotes *E. coli* virulence during mixed species CAUTI

Based on downregulation of transcripts associated with interferon regulation (*oas* and *ifi*) and monocytic chemotaxis (CCL12) during *E. faecalis*-mediated immune suppression *in vivo*, we hypothesized that suppression allows UPEC to better colonize the bladder in the presence of *E. faecalis*. To test this in a CAUTI model, we co-infected catheterized mice with 10^7^ CFU of *E. faecalis* OG1RF and 10^7^ CFU of *E. coli* UTI89, and observed no significant differences in *E. coli* titers compared to monomicrobial *E. coli* infection **(Fig 5A and 5B)**. By contrast, *E. faecalis* titers during co-infection were significantly lower in the bladders but not in kidneys, which could be a result of tissue tropism of *E. faecalis* to the kidneys or enhanced clearance as a result of the *E. coli*–driven immune activation as previously described for *E. coli*-Group B Streptococcus coinfection in the bladder **(Fig 5A and 5B)** [16, 27]. We postulated that the immunomodulatory capability of UPEC strain UTI89 may be sufficient to cause high titer CAUTI such that *E. faecalis* cannot further augment infection [28]. Therefore, we hypothesized that colonization by a non-pathogenic, commensal-like *E. coli* strain such as K12 strain MG1655, deficient for LPS O-antigen expression, may be enhanced by *E. faecalis*-mediated immune modulation [29]. To test this, we infected catheterized mice with 10^7^ *E. coli* K12 strain MG1655 alone, or 10^7^ each of *E. coli* together with *E. faecalis* at equal ratios. Similarly to infection with UTI89, *E. coli* CFU were not different following co-infection with *E. faecalis* in the bladder at 24 hpi compared to *E. coli* mono-species infection, and *E. faecalis* CFU were significantly decreased **(Fig 5C)**. By contrast, *E. coli* CFU were significantly increased in the kidneys following co-infection with *E. faecalis*, while *E. faecalis* CFU were unchanged **(Fig 5D)**. Collectively, these infection studies show that the presence of immune-modulatory organisms such as *E. faecalis*, in the context of a polymicrobial CAUTI, can increase the pathogenicity of otherwise non-virulent infectious organism and increase host vulnerability to infection by otherwise commensal organisms.

**Fig 5.**
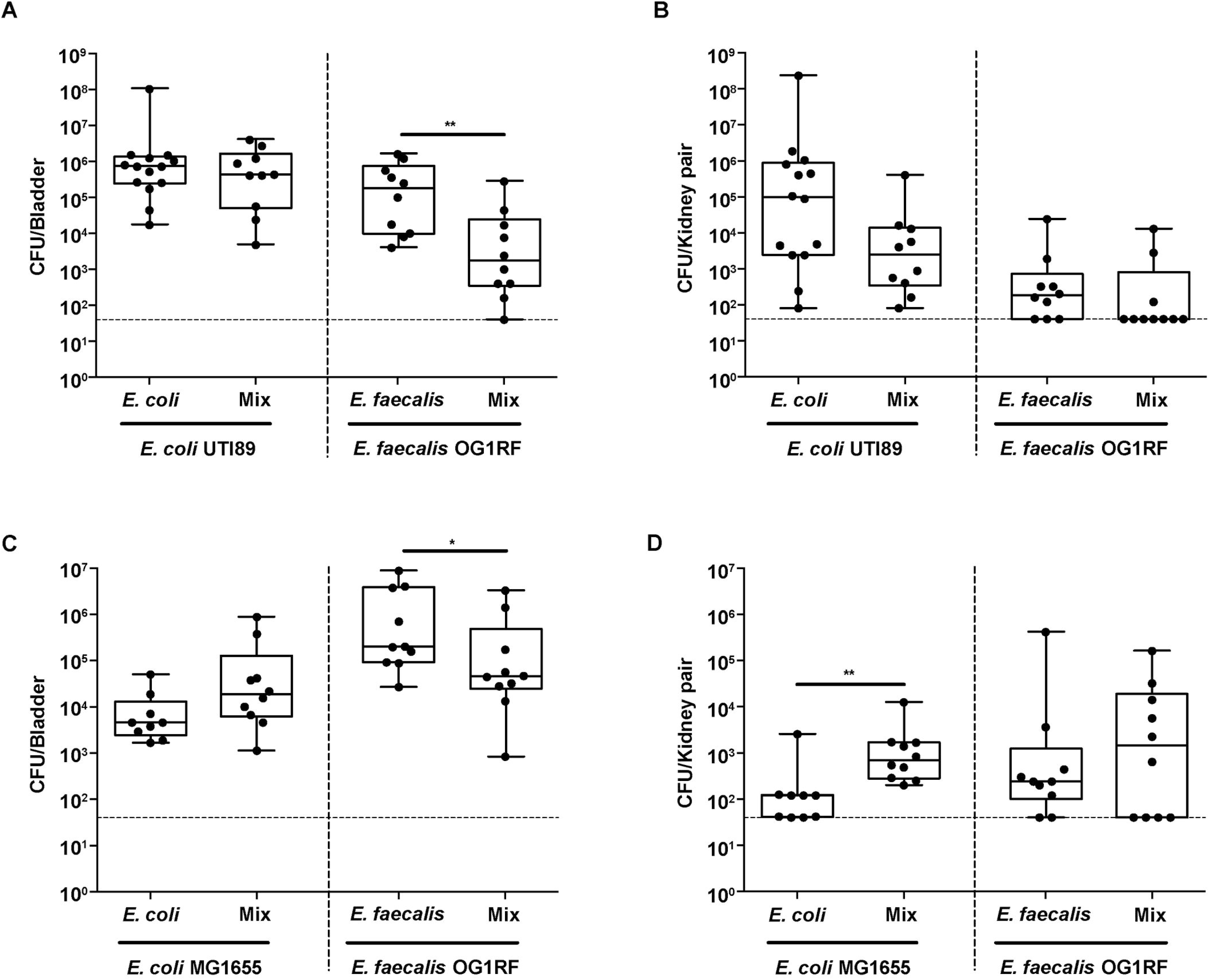
*E. faecalis* promotes *E. coli* MG1655 infection during mixed species CAUTI *in vivo*. Female C57BL/6NTac mice were implanted with 5 mm silicon catheters in the bladder and infected with 10^7^ CFU of *E. coli* UTI89 or MG1655 alone, 10^7^ CFU of *E. faecalis* OG1RF alone, or a 1:1 mixture of 10^7^ CFU of *E. coli* and10^7^ CFU of *E. faecalis*. (A) Bladder and (B) kidney titers from *E. coli* UTI89 and *E. faecalis* mono-and co-infection. (C) Bladder and (D) kidney titers from *E. coli* MG1655 and *E. faecalis* mono-and co-infection. After 24 hours, bladders and kidneys were removed and CFU enumerated. Data were combined from 2 independent experiments (5-7 mice per group). Boxes represent the 25th and 75th percentile with the middle line indicating the median. Whiskers represent the minimum and maximum values of the dataset. Significance was determined by the non-parametric Mann-Whitney test. \**P*<0.05, \*\**P*<0.01. The dashed horizontal line represents the limit of detection (LOD) of the assay. Titers below the LOD were assigned a value of the LOD for visualization on the log scale and 0 for statistical analyses.

## Discussion

Bacterial immunomodulatory functions can alter infection sites leading to increased susceptibility to colonization and persistence [30, 31]. *E. faecalis* can augment the immune response in a variety of cell types, including intestinal epithelial and mouse macrophage cell lines [14, 15, 30]. Recently, it was shown that *E. faecalis* strains V583 and E99 suppress NF-κB activation of intestinal epithelial cells and RAW264.7 macrophages at MOI 100 [30]. In contrast to reports of high MOI immune suppression, infection of RAW264.7 macrophages and bone marrow-derived macrophages with *E. faecalis* strain E99 at MOI 10 results in NF-κB activation [30]. These discrepant reports of NF-κB activation and suppression by *E. faecalis* underscore the need for further investigation into *E. faecalis* immunomodulatory activities within macrophages. Here, we resolve previous conflicting reports and show that both *E. faecalis* strains V583 and OG1RF prevent NF-κB activity in RAW264.7 macrophages in a dose-dependent manner.

Several *E. faecalis* virulence factors modulate immunity during infection, including aggregation substance (AS), gelatinase, and TcpF [30, 32, 33]. AS promotes phagocytosis and internalization into macrophages via interaction with complement receptor type 3. After internalization AS can resist superoxide killing leading to increased survival in macrophages [34]. In addition, gelatinase facilitates innate immune evasion by interacting with the complement system to reduce opsonization and to decrease neutrophil recruitment [32, 33, 35]. TcpF is a TIR domain-containing protein and interferes with Toll-like receptor (TLR)-MyD88 interactions, which also depend on MyD88 TIR domain-mediated interactions. As a result, *E. faecalis* TcpF expression results in decreased NF-κB p65 translocation in RAW macrophages [30, 36]. TcpF is present in *E. faecalis* V583 and is enriched in UTI isolates, but is absent in OG1RF [30, 36]. Since we observed NF-κB modulation by both *E. faecalis* OG1RF and V583, TcpF is unlikely to be the factor mediating high-level NF-κB suppression in macrophages. Instead, our data suggests that the *E. faecalis* factor, which prevents NF-κB activity, is a heat-modifiable molecule. Other Gram-positive pathogens secrete heat-modifiable immune-modulatory molecules. For example, *Staphylococcus aureus* superantigen-like proteins (SSLs) have immune modulatory functions, such as inhibiting IgA-mediated immune responses and by targeting neutrophils to limit the attachment to endothelial cells [37-42]. SSL3 can downregulate TLR2-mediated production of IL-8 by binding competitively with PAMP ligands of TLR2 [43]. Our work indicates that *E. faecalis* may possess similar secreted factors that modulate NF-κB activation and prevent bacterial clearance by host immune cells.

A large proportion of *E. faecalis* infections are polymicrobial and *E. faecalis is* frequently co-isolated with *E. coli* from urinary tract and wound infections [25, 31, 44-46]. Given the prevalence of *E. faecalis* in polymicrobial interactions, we performed *in vitro* and *in vivo* experiments to study the contribution of *E. faecalis* to co-infection outcomes. We found that *E. faecalis* prevented NF-κB activity during co-infection with live *E. coli* K12 strain MG1655 and UPEC strain UTI89 *in vitro* and augmented *E. coli* K12 strain MG1655 titers in the kidneys. Similar to our findings in this study, Gram-positive uropathogens *Staphylococcus saprophyticus* and Group B *Streptococcus* induce minimal pro-inflammatory responses in the urinary tract, and the latter limits UPEC pathogenesis in mice [27, 31, 47-49]. Taken together, our findings suggest that *E. faecalis* modulation of the immune response may promote the survival of co-infecting pathogens resulting in more severe infection.

Synergistic polymicrobial infections are increasingly recognized for their contribution to both disease severity and persistence [21, 31]. Here we show that *E. faecalis* modulated the host response and promoted infection by a co-infecting *E. coli* strain, which is otherwise non-virulent. Importantly, *E. faecalis* presence in the urinary tract, especially when titers are low, has historically been considered a commensal contaminant of questionable pathogenic significance [50]. Our findings call into question that supposition and raise the prospect that *E. faecalis* not only augments *E. coli* infections, but may also promote infection by a larger spectrum of uropathogens. Continued efforts are needed to dissect these polymicrobial molecular interactions to allow for better diagnostics and precision treatment, especially as UTI pathogens are increasingly resistant to antibiotics of last resort [51].

## Materials and Methods

### Bacterial strains and growth conditions

Uropathogenic *E. coli* (UPEC) strain UTI89 [52, 53] and *E. coli* K12 strain MG1655 [54] were grown overnight in Luria-Bertani (LB) broth or agar at 37°C under static conditions. *E. faecalis* strain OG1RF [55] or V583 [56] were grown statically in brain heart infusion (BHI) broth or agar at 37°C overnight. Overnight cultures of bacteria were centrifuged at 6,000 *g* for 5 minutes and resuspended in PBS at OD_600_ 0.7 (2 x 10^8^ CFU/ml) for *E. faecalis* and at OD_600_ 0.4 (2 x 10^8^ CFU/ml) for *E. coli*.

### Cell culture

RAW-Blue cells derived from RAW 264.7 macrophages (Invivogen), containing a plasmid encoding a secreted embryonic alkaline phosphatase (SEAP) reporter under transcriptional control of an NF-κB-inducible promoter, were cultivated in Dulbecco Modified Eagle medium containing 4500 mg/L high glucose (1X) with 4.0 nM L-glutamine, without sodium pyruvate (Gibco) and supplemented with 10% fetal bovine serum (FBS; PAA) supplemented with 200 μg/ml Zeocin at 37^°^C in 5% CO_2_.

### RAW-Blue macrophage infection

RAW-Blue cells were seeded in a 96 well plate at 100,000 cells/well in 200 μl of antibiotic-free cell culture media. Following overnight incubation, the cells were washed once with PBS and fresh media was added. The SEAP reporter assay was established by empirically defining the minimal agonist (lipopolysaccharide (LPS) or lipoteichoic acid [57]) concentration that induced the maximum SEAP activity in the absence of cell death. Cells were stimulated using LPS EB ultrapure purified from *E. coli* O111:B4 (Invivogen) (100 ng/ml) or LTA derived/purified from *Staphylococcus aureus* (Invivogen) (100 ng/ml) as positive controls, or media alone as a negative control. RAW-Blue cells were infected with *E. faecalis* (at MOI of 100:1, 50:1, 25:1, 10:1 and 1:1) for 6 hours with or without TLR agonists. Overnight bacterial cultures were centrifuged and resuspended in cell culture media. For infection experiments, live bacterial cultures were diluted to achieve the desired multiplicity of infection with macrophages (MOI). Alternatively, bacteria were heat-killed (80°C for 1 hour) prior to addition to macrophage cultures. For co-infection experiments, RAW-Blue cells were simultaneously infected with *E. coli* K12 strain MG 1655 (1:1 MOI) or *E. coli* UTI89 (MOI of 0.125:1) and *E. faecalis* OG1RF (MOIs of 100:1, 50:1, 25:1, 10:1 and 1:1). Heat-killed bacteria were verified by the absence of viable bacteria when plated on BHI agar.

### Collection of bacteria cell-free culture supernatants

Bacteria were grown in cell culture media for 6 hours, and bacteria-free culture supernatants were collected after centrifugation (6,000 *g*) followed by filtration (using a 0.2 μm syringe filter). Alternatively, supernatants were collected after infecting macrophages with bacteria at various MOIs and then filtered by using a 0.2 μm syringe filter. Sterility of bacteria-free culture supernatants were verified by the absence of viable bacteria when plated on BHI agar.

### NF-κB reporter assay

Post-infection, 20 μl of supernatant was added to 180 μl of QUANTI-Blue reagent (Invivogen) and incubated overnight at 37^°^C. SEAP levels were determined at 640 nm using a TECAN M200 microplate reader. All experiments were performed in triplicate.

### Cell viability assay

Simultaneously with supernatant collection for SEAP determination, culture supernatants were collected from each well to measure lactate dehydrogenase (LDH) release, using an LDH cytotoxicity assay (Clontech) according to manufacturer’s instructions. Background LDH activity was determined using mock (PBS) treated RAW-Blue cells. Maximal LDH activity was determined by lysing cells with 0.2% Triton-X. Each condition was carried out in triplicate. Percentage cytotoxicity was calculated as follows: (sample absorbance-background absorbance)/(maximal absorbance-background absorbance) x 100.

### Luminex MAP analysis

Supernatants were collected from RAW-Blue cells 6 hours post-infection and stored at −80°C until assessment by the Bio-Plex Pro mouse cytokine 23-plex assay kit (Bio-Rad Laboratories), according to manufacturer’s recommendations [58]. All samples were assessed using the same kit lot and at the same time to avoid inter-assay variability.

### Catheterization and bacterial infections

Mouse experiments were performed with ethical approval by the ARF-SBS/NIE Nanyang Technological University Institutional Animal Care and Use Committee under protocol ARF-SBS/NIE-A0247. Catheters were implanted into bladders of mice followed by bacterial inoculation via a transurethral catheter as previously described [18, 59]. Briefly, 6-8 week old female C57BL/6NTac mice (InVivos Pte Ltd, Singapore) were anesthetized with isoflurane (4%). Inoculum volumes of 50 μl, containing a bacterial suspension of either single or polymicrobial species prepared in PBS: (i) 10^7^ CFU of *E. coli* K12 strain MG1655 with 10^7^ CFU *E. faecalis* OG1RF, and (ii) 10^7^ CFU of *E. coli* UTI89 with 10^7^ CFU *E. faecalis* OG1RF. Single species controls (10^7^ CFU of *E. coli* K12 strain MG1655, 10^7^ CFU of *E. faecalis* OG1RF, or 10^7^ CFU of *E. coli* UTI89) were performed alongside polymicrobial infections. Animals were euthanized by carbon dioxide inhalation and cervical dislocation and bladders and kidneys were aseptically removed and homogenized in 1 ml PBS for CFU enumeration by serial dilution on MacConkey agar or BHI agar supplemented with 10 μg/ml colistin and 10 μg/ml nalidixic acid to isolate *E. coli* or *E. faecalis*, respectively. To identify bacterial species other than the inoculated *E. coli* or *E. faecalis*, serial dilutions were also plated on LB and BHI. Data are combined from 2 independent experiments (5-7 mice per group). Animals without catheters at the time of sacrifice were not included in the analyses.

### RNA-sequencing of infected bladders

Catheterized mice were infected as described above with 10^7^ *E. coli* UTI89, or mixed at a 1:1 ratio with 10^7^ *E. faecalis* OG1RF, in 50 μl PBS. After 24 hours, whole bladders were removed and incubated overnight in RNAlater (Qiagen) to allow complete tissue penetration by the protectant prior to storage at −80°C. RNA was extracted as described [60]. For each sample condition, a total of three sequencing libraries were constructed from 50-200 ng of rRNA-depleted RNA. 2 nM of each library was pooled at equal volumes and sequenced using an Illumina Hiseq®2500 v.2 (Illumina),150 bp paired-end.

### Analysis of RNA-sequencing

Sequencing data are available at NCBI’s BioProject (accession no. PRJ-NA335539). RNA sequencing results were analyzed as described in [19]. Briefly, reads were quality checked and adapters trimmed with cutadapt-1.4.1 using default parameters. The mm10 mouse genome was used as reference for tophat-2.0.11.Linux_x86_64 [61] and transcriptional read counts obtained using HTSeq-0.6.1 [62] with default parameters with a non-stranded analysis. Uness otherwise stated, all further analyses were performed in the R statistical computing environment (version 3.3.3) [63]. Differential analysis of *E. coli* mono-species infected animals to *E. coli* and *E. faecalis* OG1RF co-infected animals was performed using the R/Bioconductor package DESeq2, (version 1.40.1) [64]; using default settings from that package. The NCBI file gene2refseq (downloaded 03/03/2016) was used to convert Refseq to Entrez identifiers for further analysis of Gene Otology annotations and Immunological Genome Project (ImmGen) data for functional analysis.

### Functional analysis of bladder transcriptome

Processing of ImmGen expression data was performed in R 3.2.2 [63]. Briefly, all 681 CEL files [65] were processed using the RMA method with the R/Bioconductor package oligo (version 1.34.0). Annotation was done using the R/Bioconductor package *mogene10sttranscriptcluster.db* (version 8.4.0), with expression profiles for each immune cells referenced from Jojic *et al.* Enrichment score was calculated for each immune cell type [66], and examined differentially expressed genes in the top 1% of the score distribution for each cell type, and compared these to equal sized cohorts of randomly selected genes (S3 Table). This analysis was conducted separately in up-and down-regulated gene sets in the two–group comparison. Ontology analysis was performed using the R/Bioconductor package GO.db (version 3.4.0) and the gene2go file from the NCBI Gene database (downloaded 03/03/2016) using modified code from the R/Bioconductor package ontoTools (version 1.28.0) [67]. To test if differentially expressed genes associate with specific gene sets, we constructed 2-by-2 contingency tables and categorized genes based on whether they are differentially expressed or not for each included Gene Ontology Biological Process term. Random assignment was tested using Fisher’s Exact Test [68], and corrected for the number of terms using the Benjamini-Hochberg correction [69]. We filtered terms using their *information content (IC)*, [70], based on their frequency of occurrence between 3 and 4, resulting in a set of 157 included terms that represent an appropriate trade-off between the total number of included terms and specificity of functional insight. The entire R workflow and input data files are available online (https://github.com/rbhwilliams/Kline-polymicrobial-infection-paper).

### Statistical Analysis

Statistical analyses were performed using GraphPad Prism software (Version 6.05 for Windows, California, United States). SEAP assays were analyzed using one-way ANOVA with Tukey’s multiple comparison. Cytokine readings for Luminex MAP analysis were analyzed using Kruskal Wallis test. CFU titers were compared using the non-parametric Mann-Whitney U test. *P*-values less than 0.05 were deemed significant. Cytokine comparisons were performed using the Mann-Whitney U test and further comparison done using principal component analysis in R (version 3.3.2) with packages *factoextra* (Version 1.0.4) and FactoMineR (version 1.34). Heatmap data reflect log_2_ transformation of the raw data and are plotted in R (version 3.3.2) using the R package, pheatmap (version 1.0.8).

## Acknowledgments

We thank Krithika Arumugam for her assistance with sequence data processing. This work benefitted from data assembled by the ImmGen Consortium [26]. BYQT, HMSG, KKLC, SBT SH, RBHW, and KAK were supported by the National Research Foundation and Ministry of Education Singapore under its Research Centre of Excellence Programme. BYQT, HMSG, KKLC, SBT, SH, and KAK were supported by the National Research Foundation under its Singapore NRF Fellowship programme (NRF-NRFF2011-11) and by the Ministry of Education Singapore under its Tier 2 programme (MOE2014-T2-1-129). FG and RD were supported by Singapore Immunology Network core funding, Agency for Science, Technology and Research (A*STAR), Singapore. The funders had no role in study design, data collection and analysis, decision to publish, or preparation of the manuscript.

